# Cadmium treatment induces endoplasmic reticulum stress and unfolded protein response in *Arabidopsis thaliana*

**DOI:** 10.1101/2022.09.23.509148

**Authors:** Maria De Benedictis, Antonia Gallo, Danilo Migoni, Paride Papadia, Pietro Roversi, Angelo Santino

## Abstract

We report about the response of *Arabidopsis thaliana* to chronic and temporary Cd^2+^ stress, and the Cd^2+^ induced activation of ER stress and unfolded protein response (UPR). Cd^2+^-induced UPR proceeds mainly through the bZIP60 arm, which in turn activates relevant ER stress marker genes such as BiP3, CNX, PDI5 and ERdj3B in a concentration- (chronic stress) or time- (temporary stress) dependent manner. A more severe Cd-stress triggers programmed cell death (PCD) through the activation of the NAC089 transcription factor. Toxic effects of Cd^2+^ exposure are reduced in the *Atbzip28/bzip60* double mutant in terms of primary root length and fresh shoot weight, likely due to reduced UPR and PCD activation. We also hypothesised that the enhanced Cd^2+^ resistance of the *Atbzip28/bzip60* double mutant is due to an increase in brassinosteroids signaling, since the amount of the brassinosteroid insensitive1 receptor (BRI1) protein decreases under Cd^2+^ stress only in Wt plants. These data highlight the complexity of the UPR pathway, since the ER stress response is strictly related to the type of the treatment applied and the multifaceted connections of ER signaling. The reduced sensing of Cd^2+^ stress in plants with UPR defects can be used as a novel strategy for phytoremediation.

## Introduction

Cadmium (Cd) is a non-essential metal and it is highly toxic. It is well known that Cd-contaminated sites are a serious worldwide concern (Khan et al., 2021) due to the toxic impact of Cd on human health (Suhani et al., 2021) and environmental ecosystems (Hayat et al., 2019). As well as inducing tumours, Cd also damages liver, kidneys, bones and heart (Nordberg et al., 2018; Genchi et al., 2020).

In plants, Cd^2+^ initially induces chlorosis and reduce growth (Jali et al., 2016), due to inhibition of photosynthetic activity, chlorophyll biosynthesis and interference with the uptake of essential metals. As well as phenotypic alterations, Cd^2+^ also induces the production of reactive oxygen species (ROS) with consequent lipid peroxidation of membranes and inhibition of antioxidative enzymes (Haider et al., 2021). Transcriptomic (Herbette et al., 2006; Baliardini et al., 2016), proteomic and metabolomic studies (Sarry et al. 2006; De Benedictis et al., 2018) have highlighted the effects of Cd^2+^ on primary and secondary metabolism. However, strategies to improve plant tolerance to Cd^2+^ based on the overexpression of genes involved in metal detoxification or metal transport have so far achieved limited success (Thomine et al., 2000; Lee et al., 2003; Li et al., 2005; Cailliatte et al., 2009; Haydon et al., 2012).

Little attention has been directed to the effects of Cd accumulation in plant tissues and Endoplasmic Reticulum (ER) stress, glycoprotein folding Quality Control (ERQC) and ER associated glycoprotein degradation (ERAD).

Glycoproteins folding relies on a finely tuned balance between ERQC and ERAD. ERQC surveys and assists glycoprotein folding, whereas ERAD flags terminally misfolded glycoproteins for degradation *via* their demannosylation, retrotranslocation and proteasome associated degradation. Both pathways use N-glycans as markers to label the folding status of glycoproteins. In the case of ERQC, the presence of a glucose moiety on a high-mannose glycan (Glc_1_Man_9_GlcNAc_2_) promotes protein association with ER chaperones of the calnexin cycle, i.e. the lectins calnexin (CNX) and calreticulin (CRT), lectin-associated chaperones (e.g. BiP3) and protein disulfide isomerases (*e*.*g*. PDI5) – which in turn assist the folding of the monoglucosylated glycoprotein (Kornfeld and Kornfeld, 1985; Vitale, 2001; Strasser, 2016). Monoglucosylated glycoproteins are generated either by ER α-Glucosidase II (α-Glu II) acting on a glycoprotein carrying a di-glucoslyated glycan (Glc_2_Man_9_GlcNAc_2_, the glucoses originating from the dolichol N-glycan precursor); or by UDP-Glucose Glycoprotein Glucosyltransferase (UGGT), which transfers Glc from UDP-Glc to a Man_9_GlcNAc_2_ glycan on a misfolded glycoprotein. Once a glycoprotein has reached the correct folding, it can proceed to the Golgi and from here move down the rest of the secretory pathway. Glycoproteins that are not able to reach their correct fold are eventually degraded by the ERAD pathway through the ER degradation associated mannosidases (EDEMs) (Brodsky and McCracken, 1999).

Several physiological conditions in plants, such as seed/pollen maturation (Vitale and Boston, 2008), (a)biotic stresses (heat/salt stress, bacterial/virus infections) or the treatment with chemical stressors (tunicamycin, the proline analog azetidine-2-carboxylase or dithiothreitol) (Iwata and Koizumi, 2005a) are characterised by a high rate of glycoprotein biosynthesis. All these conditions have the potential to increase the accumulation of misfolded glycoproteins in the ER and alter the effectiveness of ERQC/ERAD in coping with ER stress. Under ER stress, the Unfolded Protein Response (UPR) is activated to restore glycoprotein homeostasis.

The plant UPR pathway has been only recently elucidated. Similarly to UPR in animals (Li and Howell, 2021), there are two UPR arms in plants. One arm involves the ER-localised inositol requiring enzyme 1 (IRE1) (Deng et al., 2011; Nagashima et al., 2011), which is involved in the splicing of the mRNA encoding bZIP60 (bZIP60u) transcription factor (TF) to generate an isoform lacking the transmembrane domain, bZIP60s (Li and Howell, 2021), which is then translated and translocated into the nucleus. The other UPR arm in plants is mediated by two ER-resident TFs, namely bZIP17 and bZIP28, which upon ER stress, are translocated to the Golgi and cleaved by the Golgi resident subtilisin-like serine proteases S1P and S2P, respectively, causing release and translocation to the nucleus of the b-ZIP cytosolic domain (Liu et al., 2007a; Iwata et al., 2017). Once in the nucleus, bZIP60s and bZIP17/bZIP28 converge in regulating the expression of stress response genes. As a result, glycoprotein transcription is reduced and glycoprotein folding capacity increased (*via* the induction of chaperones, e.g. CNX, BiP3), lowering the level of unfolded glycoproteins in the ER (Deng et al., 2013; Liu and Howell, 2016). If ER glycoproteostasis is not restored, UPR can activate programmed cell death (PCD) (Iwata and Koizumi, 2005b; Xu et al., 2013) through the nuclear translocation of other transcription factors, e.g. NAC089 (Yang et al., 2014).

The *Arabidopsis thaliana* (*A. thaliana*) *bzip28/bzip60* double mutant is highly susceptible to tunicamycin (Tm) and displays a significant reduction in primary root and shoot growth (Lai et al., 2018; Ruberti et al., 2018) as well as vulnerability to necrotrophic fungi, e.g. *Drechslera gigantean* (Samperna et al., 2021). Conversely, stress tolerance increases in transgenic plants overexpressing the active forms of the bZIP transcription factors (Fujita et al., 2007; Tang et al., 2012; Zhang et al., 2015; Liu et al., 2008).

Intriguingly, the double mutant *bzip28/bzip60* is more tolerant to intense light (Beaugelin et al., 2020) and chronic Cd^2+^ exposure (Xi et al., 2016), suggesting a complex regulation of UPR pathways based on the type of stress applied. The present work aims at studying Cd-induced ER stress at the physiological and molecular levels. To do this, we characterised the effects of Cd^2+^ on Wt and the *bzip28/bzip60* double mutant by studying: i) chronic stress induced by germinating and growing plants in the presence of two Cd^2+^ concentrations (25 and 50 μM); ii) strong temporary stress (150 μM Cd^2+^) imposed for different durations (24, 48 or 72 h).

We report for the first time about the impact of these two types of stresses on the transcription of key genes of the UPR, ERQC/ERAD and PCD pathways, the overall *N*-glycosylation process and glycoprotein folding capacity in *A. thaliana*.

## Materials and methods

### Plant lines, growth conditions and stress assays

*A. thaliana* Wild type (Wt) ecotype Columbia 0 (Col-0), the double mutant *bzip28/bzip60* (Ruberti et al., 2018) and *bri1-9* mutant (Vert et al., 2005) were grown in one-half strength Murashige and Skoog (MS) medium at 22°C with 60% relative humidity under 14 h of daylight.

For chronic stress assays, seedlings were germinated and grown for 10 days in the presence of different CdCl_2_ concentrations (0, 25, 50, 75 μM); for temporary stress assays, 10 days old plants were treated with 0 or 150 μM CdCl_2_ for 24, 48, 72 h.

Phenotypic analysis of the relative root and shoot growth rates was carried out, as reported in Meng et al. (2017).

### RNA extraction and gene expression analysis

Total RNA extraction from *A. thaliana* tissues was done using the RNeasy Plant Mini Kit (Qiagen) according to the manufacturer’s instructions. Samples were then treated with Deoxyribonuclease I and 1 μg of total RNA was used for cDNA synthesis using the SuperScript III First-Strand synthesis system kit (Invitrogen).

Expression analysis on a set of different genes involved in UPR/ERQC/ERAD/PCD (see Supplementary Table S1) was performed with the SYBR Green Supermix (Bio-Rad) on StepOnePlus Real-Time PCR System (Applied Biosystems) using as a template 2 μL of 10-fold dilution of synthesized cDNA. The qRT-primers efficiency, listed in Supplementary Table S2, was estimated from linear regressions of the standard curves and the reaction mix was used as indicated: 95 °C for 2 min, followed by 40 cycles of 15 s at 95 °C and 1 min at 60 °C. The specificity of the PCR amplification was confirmed by dissociation curve analyses. Values are averages from three biological replicates, performed in triplicate.

### Protein extraction and immunoblotting

Total protein was extracted from 0.15 g of plant tissues ground to powder with liquid N_2_ and suspended in 100 μl 2X SDS-PAGE sample buffer under reducing conditions. Samples were heated to 100 °C for 5 mins and centrifuged at 10,000 g for 10 mins at 4° C. Supernatants were loaded and run on a 10% SDS PAGE polyacrylamide gel, then transferred onto a PVDF membrane and blotted with anti-BRI1 (1:1000 dilution) primary antibody (Agrisera, AS121859). Densitometric quantification of BRI1 bands was done using the stain-free normalisation technique with Image Lab Software (Biorad).

For N-glycoproteins detection, the same total protein extracts were incubated with Endoglycosidase H at 37 °C for 3 h (Sigma-Aldrich) according to the manufacturer’s instructions. Both Endo H-digested samples or not were resolved on a 10% SDS-PAGE gel under reducing conditions. After transfer to a PVDF membrane, the glycoprotein bands were probed with 1 μg/ml Concanavalin A (Con A) peroxidase conjugate (Sigma-Aldrich, L6397). Signals were detected by electrochemiluminescence (Bio-Rad).

### Elemental analysis

Wt and *bzip28/bzip60* seedlings were grown for 14 days with 0, 25, and 50 μM CdCl_2_ and harvested according to Zhai et al., 2014 before analysis of metal content.

The concentration of Cd^2+^ was measured in plant samples using Inductively Coupled Plasma Atomic Emission Spectroscopy (ICP-AES, ThermoFisher Scientific, USA, Waltham, Massachusetts). Each sample was weighted and mixed with 4 mL of H_2_O_2_ and 6 mL of suprapure HNO_3_ 69 %, then treated at 180°C for 10 min, using a microwave digestion system (Milestone START D). The samples were then cooled, diluted with ultrapure water to a final volume of 20 mL, filtered through syringe filters (pore size 0.45 μm), and then measured for element content using an ICP-AES (Thermo Scientific, iCap 6000 Series) spectrometer. An ICP Multi-Element Standard Solution VIII (Merck, Darmstadt, Germany), consisting of 29 elements in dilute nitric acid, was used to construct the calibration lines for the spectrometer. For quantitative analysis, five standards were prepared for a calibration line covering a large range of concentrations. The calibration line had a linear correlation coefficient (r) greater than 0.99, the detection limit of the element corresponds to the lowest concentration of the calibration line. Results for each plant were expressed as the average of three different measurements, with elemental concentrations expressed as ppm (mg/kg of dry weight). The Certified Reference Material used for quality control and quality assurance was the CPAchem (Stara Zagora, Bulgaria). Multi-Element Standard Solution. The total amount of elements accumulated in the seedlings, was calculated by multiplying the element concentration by the total tissues weight of each plant.

### Statistical analysis

Statistical significance analysis used a Student’s two-tailed *t*-test with equal variance.

## Results

### A. thaliana response to chronic or temporary Cd^2+^ stress

We first carried out a phenotypic analysis of Wt plants challenged with Cd^2+^ chronic stress (25 and 50 μM CdCl_2_, for 10 days). Results in Fig. 1 indicate that Cd^2+^ stress induced a significant and concentration-dependent reduction in root length and plant biomass weight. Next, we monitored the expression profiles of the transcription factors (TFs) bZIP28 and bZIP60, the main components of the UPR pathway. In particular, bZIP60 was studied as unspliced (*bZI60u*) and spliced (*bZIP60s*) transcripts, the latter deriving from the IRE1-mediated unconventional splicing of the *bZIP60* gene.

**Fig. 1.**
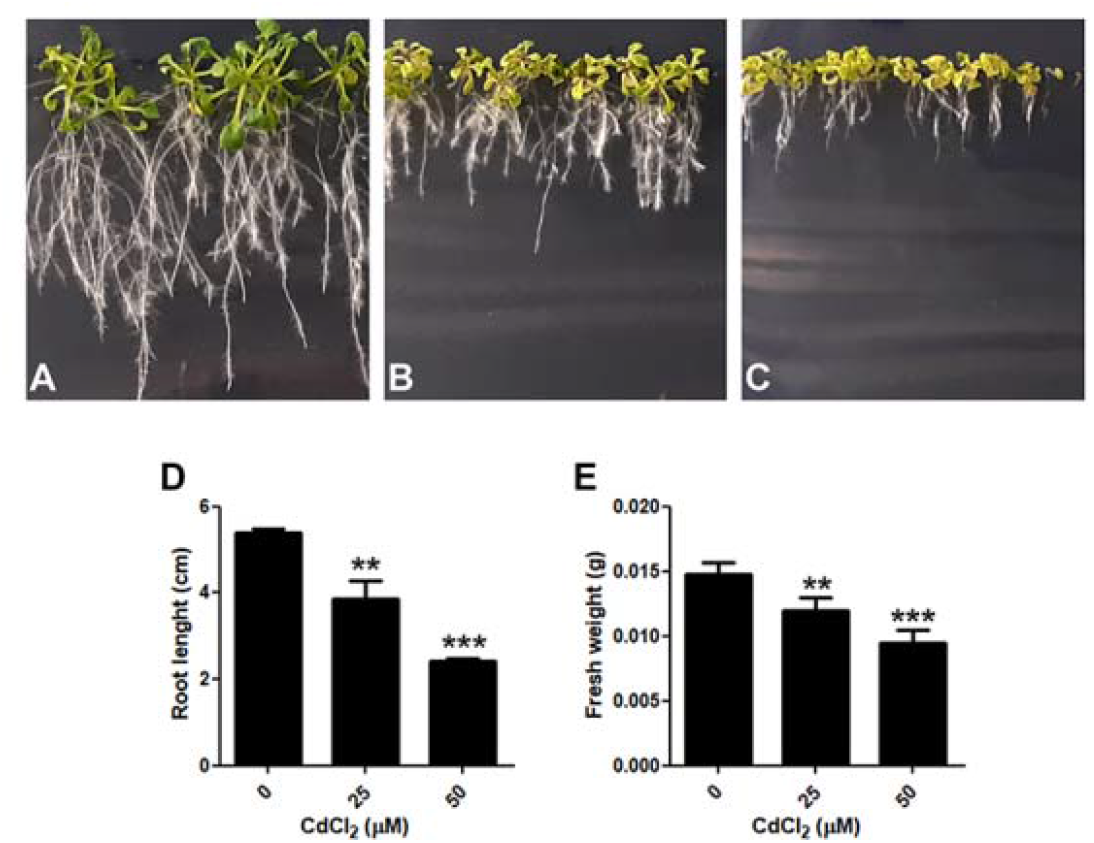
Cd-stress influences negatively the growth of *A. thaliana* Wt plants. Seedlings were grown in presence of 0 (A), 25 (B), 50 (C) μM CdCl_2_ for 10 days. Root length (D) and fresh weight (E) were measured in all conditions tested. Error bars indicate standard deviation among at least ten samples. Asterisks convey the statistical significance of the differences to control (* p ≤ 0.05; ** p ≤ 0.01; *** p ≤ 0.001).

In agreement with a previous report (Xi et al., 2016), chronic Cd^2+^ stress induced a significant upregulation of both isoforms of *bZIP60* already at 25 μM (Fig. 2A, 2B), whereas a 1.5 fold upregulation of *bZIP28* was recorded at 50 μM (Fig. 2C).

**Fig. 2.**
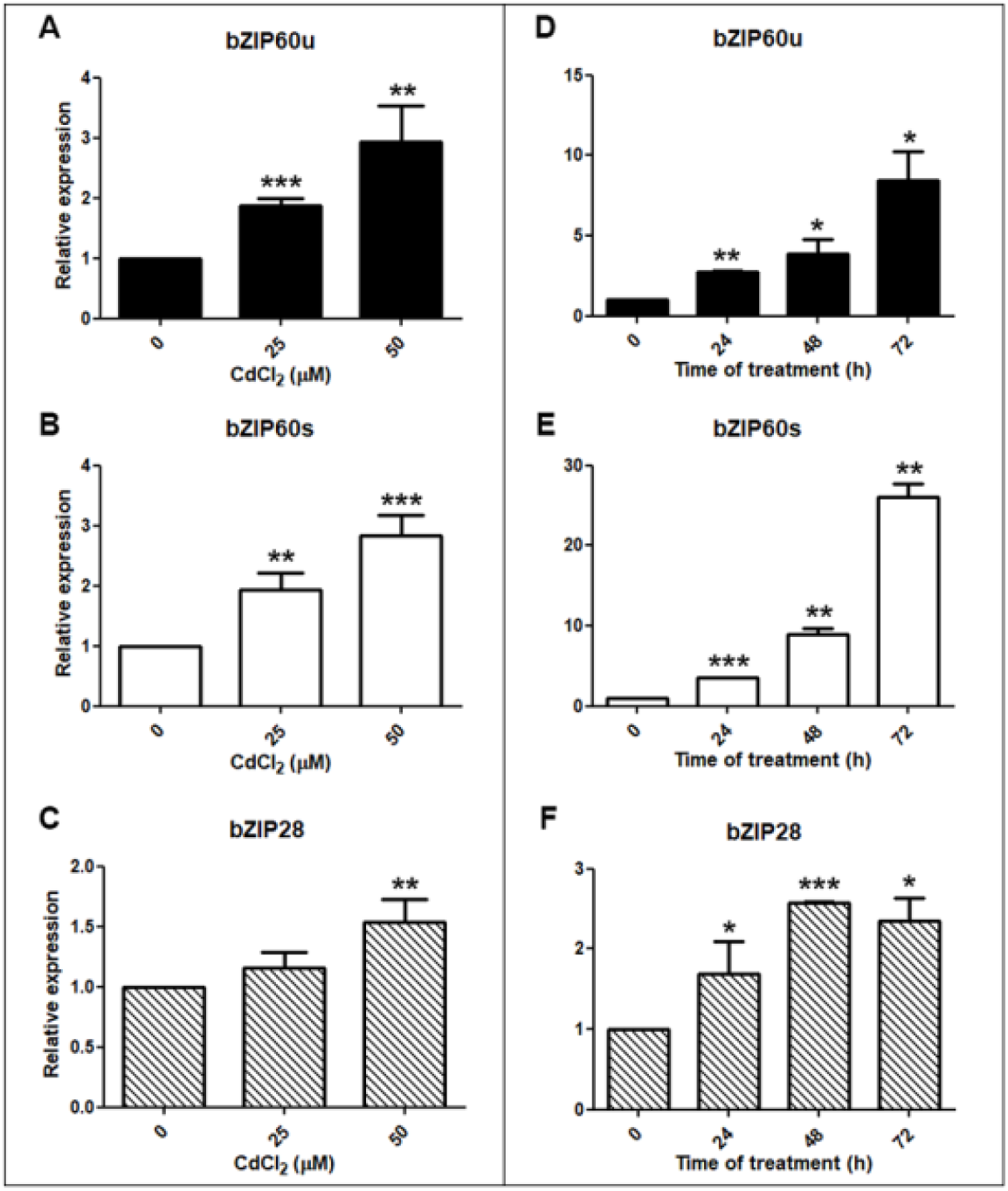
Relative expression of the unspliced *(bZIP60u*) and spliced (*bZIP60s*) forms of *AtbZIP60* and *AtbZIP28* transcription factors under Cd stress in Wt *A. thaliana*. (A-B-C) Chronic stress was induced by growing plants for 10-days with different concentrations of CdCl_2_. (D-E-F) Temporary stress was applied to 10-day-old seedlings with 150 μM CdCl_2_ at different time points. All values are normalised to Actin2 and Ubiquitin5 expression and are reported relative to Wt control (0 μM Cd) set at 1. Error bars indicates SD and asterisks indicate a statistically significant difference compared to control (* p ≤ 0.05; ** p ≤ 0.01; *** p ≤ 0.001).

Temporary Cd^2+^ stress (150 μM) induced the upregulation of all the transcripts already from 24 h after stress onset (Fig. 2 D-F), with the *bZIP60s* transcripts showing the highest activation at longer time points. In particular, *bZIP60s* was upregulated more than 20 times after 72 h compared to unstressed plants (Fig. 2E) and it was activated more strongly under severe temporary stress than chronic milder stress (Fig. 2B, 2E).

Weaker upregulation was observed for *bZIP28*, with a maximum (about 2.5 times compared to control plants) at 48 h after stress onset (Fig. 2F).

### Response of A. thaliana bzip28/bzip60 double mutant to Cd^2+^ stress

3 days old *bzip28/bzip60* double mutant plantlets were reported to display enhanced tolerance to chronic Cd^2+^ stress (up to 75 μM concentration for 2 weeks) (Xi et al.; 2016). We verified the phenotypic response of the *bzip28/bzip60* double mutant to chronic Cd-stress induced by three Cd^2+^ concentrations (25, 50 and 75 μM of CdCl_2_, for 10 days). Images and histograms in Fig. 3 confirm a higher tolerance of this mutant to Cd^2+^ stress. In comparison with Wt plants, a significant increase in primary root length was recorded at all the tested concentrations in *bzip28/bzip60*. Plant biomass was also significantly higher in the double mutant with respect to Wt at 50 and 75 μM Cd^2+^ (Fig. 3E, F). This enhanced Cd resistance of the double mutant is not due to a lower Cd^2+^ intake since the quantification of metal content showed no difference between the two plant lines in all conditions tested (Fig. S1).

**Fig. 3.**
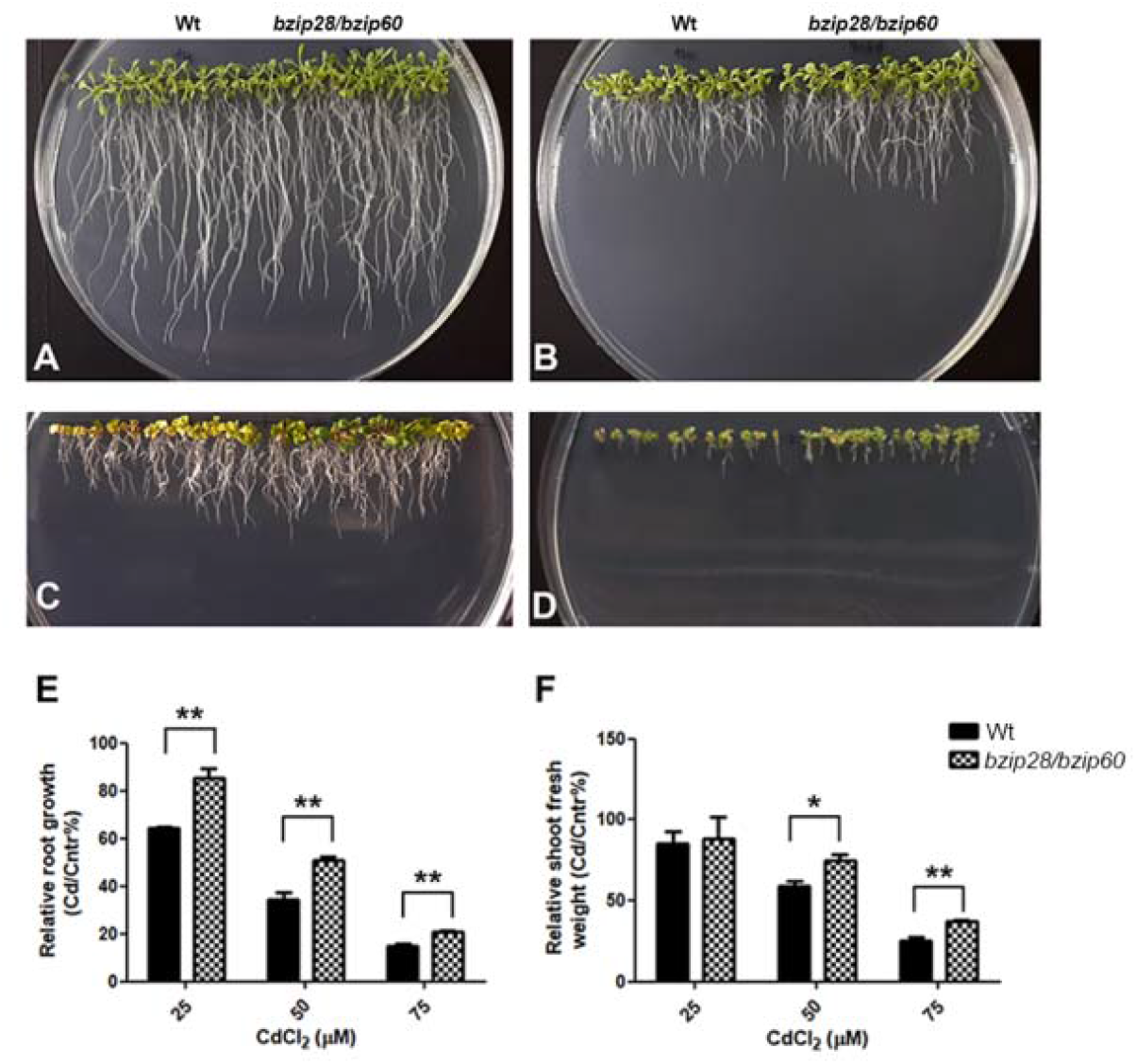
The *A. thaliana bzip28/bzip60* double mutant shows increased tolerance to CdCl_2_ stress. Wt and *bzip28/bzip60* plants treated with 0 (A), 25 (B), 50 (C), 75 (D) μM CdCl_2_ for 10 days. (E) Relative root length and (F) shoot weight of Cd-treated seedlings in comparison with the control condition. Error bars indicate standard deviation, and asterisks indicate the statistical significance of the difference between the two plant lines for each condition. The values represent the average of at least three biological replicates (* p ≤ 0.05; ** p ≤ 0.01; *** p ≤ 0.001).

In addition, we monitored the expression levels of the main markers of ER stress and UPR pathways in both Wt and *bzip28/bzip60*, when challenged with chronic or temporary Cd^2+^ stress. Under chronic treatment, the ER stress marker genes *CNX, PDI5, BiP3* and the UPR marker gene *ERdj3B* are all significantly upregulated in Wt plants already at 25 μM Cd^2+^ and their expression levels increase with increasing Cd^2+^ concentration (Fig. 4A-D).

**Fig. 4.**
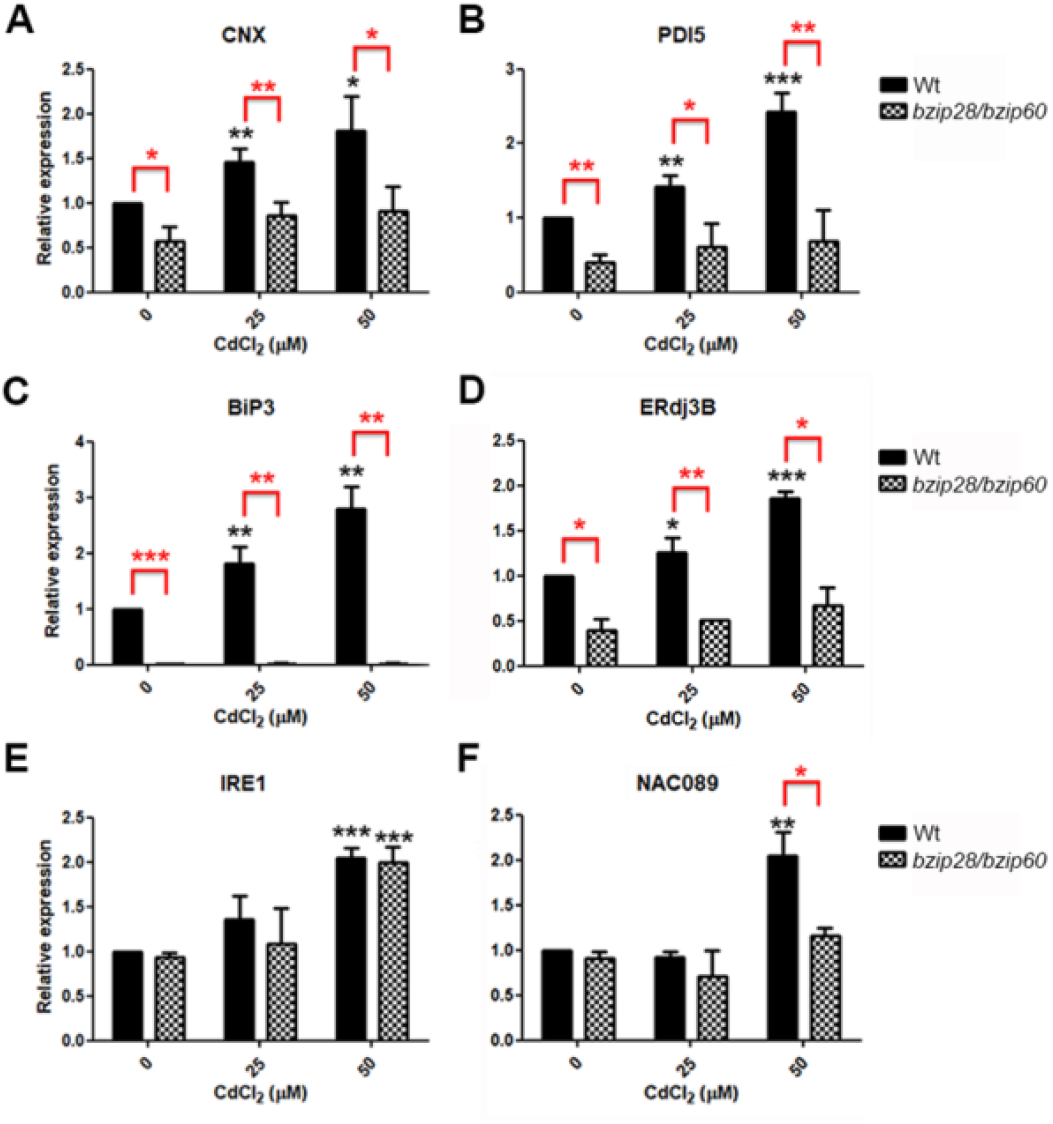
Response to CdCl_2_ induced ER stress and UPR is compromised in *bzip28/bzip60* double mutant. (A-B-C) qRT-PCR analysis of principal ER stress marker genes, *AtCNX, AtPDI5, AtBIP3*; (D-E) qRT-PCR analysis of genes involved in UPR pathway, *AtERdj3B, AtIRE1*; (F) qRT-PCR analysis of the expression level of the transcription factor involved in PCD activation, *AtNAC089*. All values are normalised to Actin2 and Ubiquitin5 expressions and are shown relative to Wt control (0 μM Cd) set at 1. Error bars represent SD. Black asterisks indicate a significant difference between control and CdCl_2_ treated seedlings of the same genotype. Red asterisks show a significant difference between the two genetic backgrounds under the same treatment conditions (* p ≤ 0.05; ** p ≤ 0.01; *** p ≤ 0.001).

Differently from Wt plants, the *bzip28/bzip60* mutant shows significantly lower transcription levels of these genes, both with no Cd treatment and upon increasing Cd^2+^ concentration (Fig. 4A-D). A similar trend is observed for the *NAC089* TF, which is significantly upregulated in Wt plants challenged with harsh Cd^2+^ stress (50 μM), but not in *bzip28/bzip60* (Fig. 4F).The levels of *IRE1* transcripts show a similar trend in Wt and *bzip28/bzip60* with a significant increase at 50 μM Cd^2+^.

Differences in the response time of the Cd-induced upregulation of the tested genes are observed when a temporary stronger Cd^2+^ stress is imposed (Fig. 5). In Wt plants, *CNX, PDI5* and *ERdj3B* genes all appear upregulated only with the most prolonged treatment (72 h), whereas *BiP3* is upregulated already with a 24 h treatment. As already shown under chronic stress, in the *bzip28/bzip60* mutant the expression levels of *CNX, PDI5* and *ERdj3B* are significantly lower than those recorded in Wt plants and the double mutant plants do not respond to Cd^2+^ stress by inducing transcription of these genes. Surprisingly, a slight but significant increase in the transcript levels of *BiP3* is recorded after 48 and 72 h in *bzip28/bzip60*. These results suggest that *BiP3* transcription is not completely under the control of *bZIP28* or *bZIP60*. Similar results are recorded for *NAC089*, whose levels show a slight but significant increase in the *bzip28/bzip60* background only after 72 h of treatment, although at lower levels compared to Wt stressed plants (Fig. 5F). No differences in the levels of *IRE1* are observed between Wt and *bzip28/bzip60* plants (Fig. 5E).

**Fig. 5.**
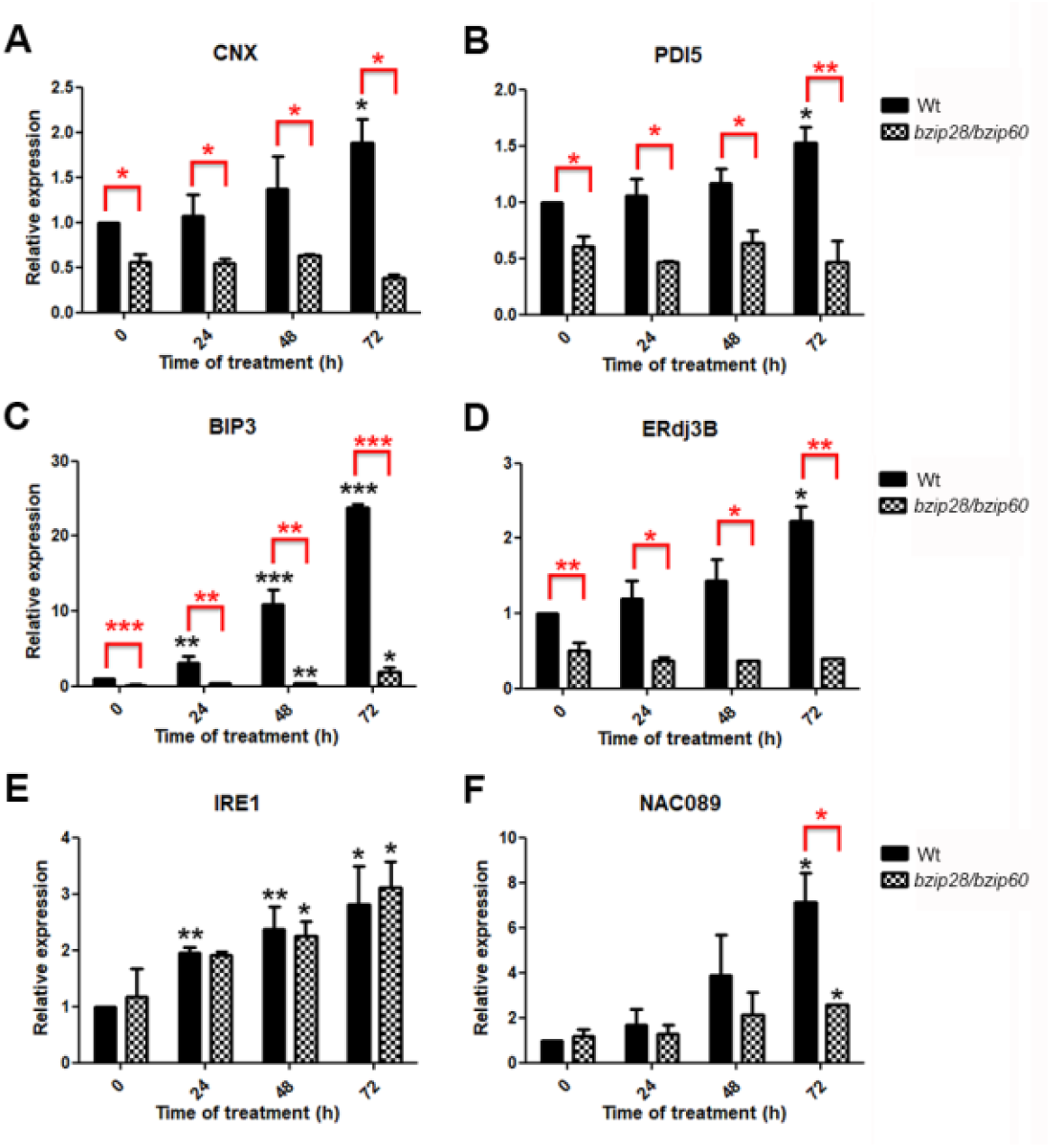
ERAD and UPR pathways are compromised in *bzip28/bzip60* double mutant both in the control condition and in the presence of 150 μM CdCl_2_ up to 72 hours of treatment. (A-B-C) qRT-PCR analysis of principal ER stress marker genes, *AtCNX, AtPDI5,AtBIP3*; (D-E) qRT-PCR analysis of genes involved in UPR pathway, *AtERdj3B, AtIRE1*; (F) qRT-PCR analysis of the expression level of the transcription factor involved in PCD activation, *AtNAC089*. All values are normalised to Actin2 and Ubiquitin5 expressions and are shown relative to Wt control (0 μM Cd), which was set to 1. Error bars represent SD. Black asterisks indicate a significant difference between control and CdCl_2_ treated seedlings of the same genotype. Red asterisks show a significant difference between the two genetic backgrounds in the same condition of growth (* p ≤ 0.05; ** p ≤ 0.01; *** p ≤ 0.001).

### Cd^2+^ stress impact on BRI1 protein levels

The brassinosteroid receptor BRI1 folds under the control of ERQC and is a key regulator of plant growth (Nagashima et al., 2018). The *bri1-9* mutant cannot fold properly, it is retained in the ER and finally degraded, with the consequence that the plant carrying this mutation is insensible to brassinosteroids and has a typical dwarf phenotype (Noguchi et al., 1999).

We checked the transcript/protein levels of this gene in the Wt and *bzip28/bzip60* mutant plants to evaluate if Cd-induced ER stress impacts on BRI1 folding capacity and this could at least in part explain the phenotypic response of *A. thaliana* showed in Fig. 1 and Fig. 3. The transcript levels of *BRI1* do not vary in Wt and *bzip28/bzip60* upon chronic Cd^2+^ stress (50 μM for 10 days) (Fig. 6A). Western blot analysis, carried out using a specific antibody raised against the BRI1 glycoprotein, shows similar protein levels in normal growth conditions in Wt and *bzip28/bzip60* plants (Fig. 6B, C). The amount of BRI1 decreases under Cd^2+^ stress in Wt plants, but not in the double mutant (Fig. 6B,C; Fig. S2). In accordance with these biochemical results, Wt plants grown in the presence of 50 μM Cd^2+^ for 4 weeks show a phenotype similar to the well known dwarf phenotype of the *bri1-9* mutant (Fig. 6D). In the case of *bzip28/bzip60* mutant, a less drastic phenotype is observed (Fig. 6 D).

**Fig. 6.**
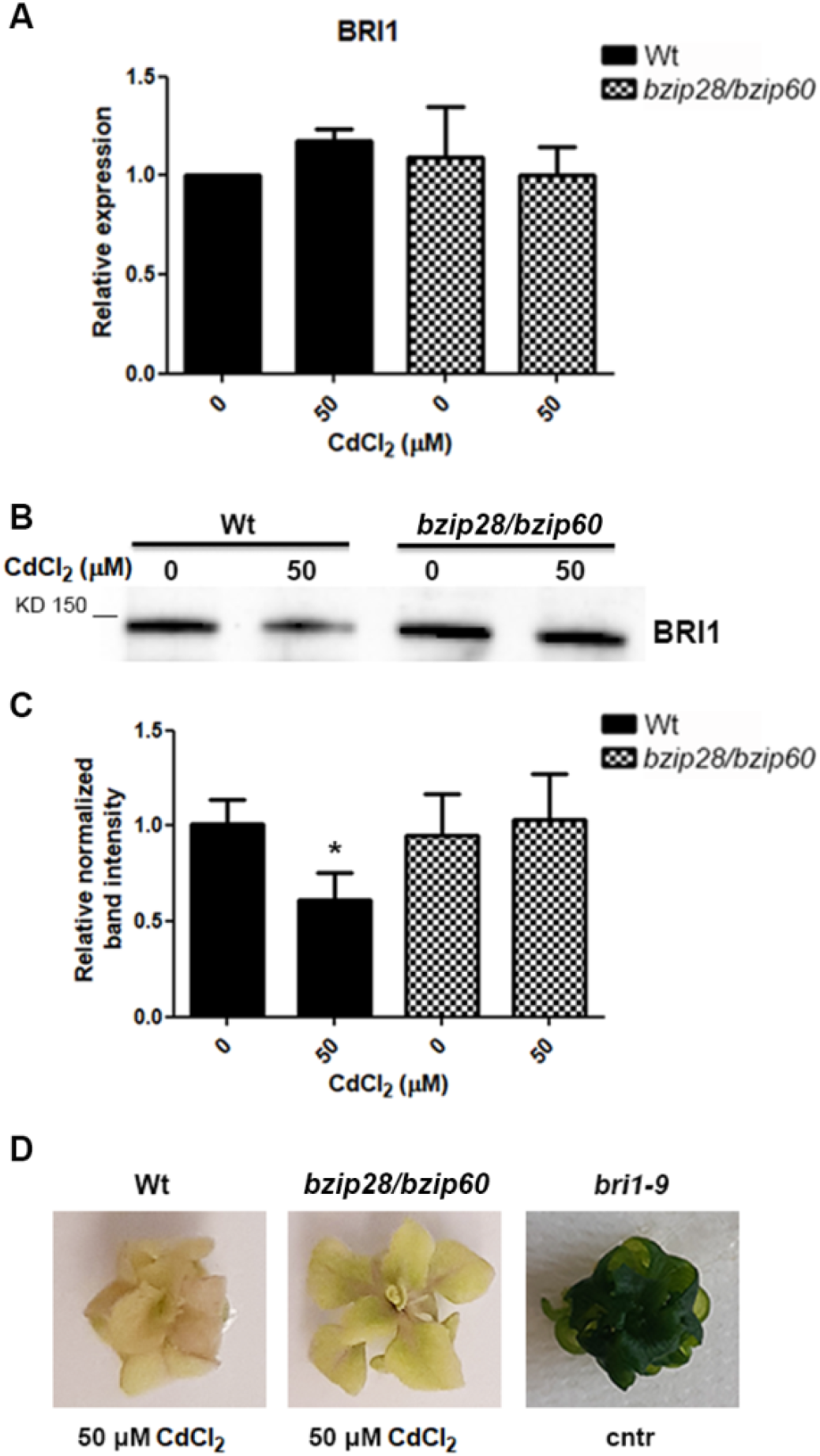
CdCl_2_ treatment impacts the protein level of brassinosteroid receptor BRI1. Seedlings of *A. thaliana* Wt and *bzip28/bzip60* were grown for 10 days with or without 50 μM CdCl_2_ and tissues were used to measure the levels of (A) transcripts by qRT-PCR and (B, C) protein by Western blotting of BRI1. The BRI1 band intensity is normalised on total proteins loaded. The reference gel lane against which all other lanes are compared is Wt control (0 μM CdCl_2_). (D) Phenotype of Wt and *bzip28/bzip60* plants grown for 4 weeks with 50 μM CdCl_2_ and *bri1-9* mutant in the regular condition of growth. qRT value is normalised to Actin2 and Ubiquitin5 expressions and is shown relative to Wt control (0 μM Cd), which was set to 1. Data are shown as means ± standard deviation. The asterisks indicate a statistically significant difference compared to Wt 0 μM CdCl_2_ (* p ≤ 0.05).

### Cd^2+^ has a minor impact on overall glycoprotein folding capacity

We explored the impact of Cd^2+^ stress on the overall glycoproteins folding capacity of *A. thaliana*. With this aim, we first quantified the transcription levels of genes encoding (ERQC) key enzymes, such as ER α-Glu II and UGGT, and the ERAD mannosidase 4 (MNS4), in Wt and *bzip28/bzip60* plants. It is of note that in *bzip28/bzip60* mutant plants, all the three ERQC genes are significantly downregulated in the control condition and under low Cd^2+^ stress (25 μM) in comparison with Wt plants (Fig. 7 A-C). However, stronger Cd^2+^ stress resulted in a significant increase in the level of *UGGT* and *MNS4*, but not of α-Glu II (Fig. 7 A-C). These results indicate the possibility that the transcription of these genes might be at least in part directly or indirectly controlled by *bZIP28/bZIP60*.

**Fig. 7.**
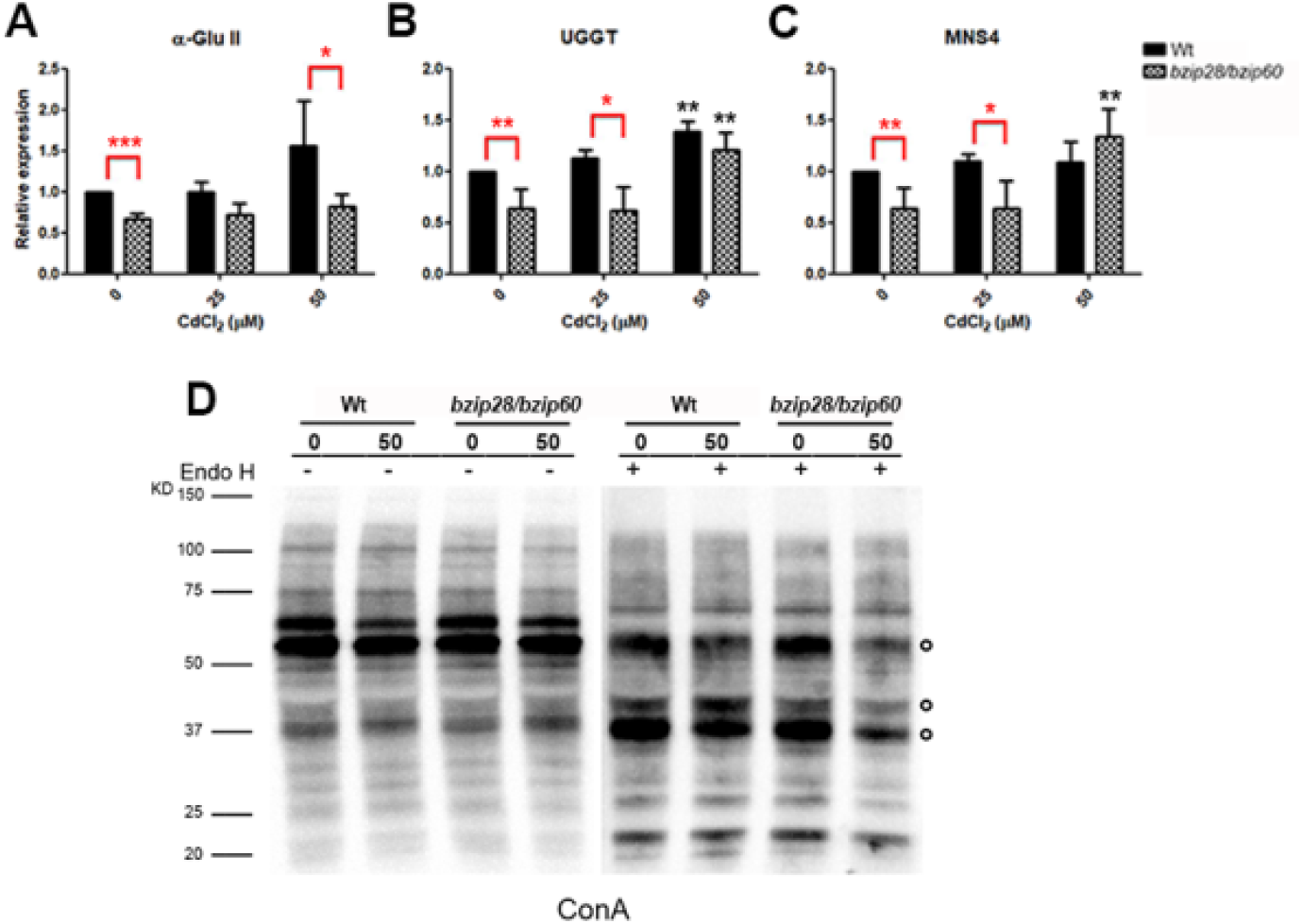
CdCl_2_ stress affects protein *N*-glycosylation only in *bzip28/bzip60* double mutant plants. (A-B) qRT-PCR analysis of transcripts of key enzymes involved in the modification of *N*-glycan chain, *At*_α_*-GluII* and *AtUGGT*; (C) qRT-PCR analysis of the transcription level of *AtMNS4*, a mannosidase responsible for directing misfolded proteins to degradation. All values are normalised to Actin2 and Ubiquitin5 expressions and are shown relative to Wt control (0 μM Cd), which was set to 1. Error bars represent SD. (D) Concanavalin A (ConA) blot assay of Endo H digestion from Wt and *bzip28/bzip60* total proteins extracted from tissues grown with or without 50 μM CdCl_2_ for 10 days. Open circles indicate differences in glycoproteins accumulation.Black asterisks indicate a significant difference between control and CdCl_2_ treated seedlings of the same genotype. Red asterisks show a significant difference between the two genetic backgrounds in the same condition of growth (* p ≤ 0.05; ** p ≤ 0.01; *** p ≤ 0.001).

In order to verify if Cd^2+^ stress impacts the overall glycoproteins pattern, plant glycoproteins, digested or not with Endoglycosidase H (EndoH), from Wt and *bzip28/bzip60* plants unstressed or challenged with 50 μM Cd^2+^ were separated on SDS-PAGE and immunoblotted with HRP-conjugated Concanavalin A (ConA). ConA is a lectin which selectively binds high mannose N-glycans of glycoproteins and is commonly used to probe ER-retained glycoproteins (Faye and Chrispeels, 1985). Cd^2+^ treatment has a slight, if any, impact on ER-retained glycoproteins (Fig. 7D; Fig. S3). Endo H digestion of the same glycoprotein extracts before ConA immunoblotting highlights the presence of some weaker glycoprotein signals in the *bzip28/bzip60* double mutant (Fig. 7D, indicated by an open circle), thus suggesting that in this genetic background some glycoproteins fold more slowly or are retained in the ER for longer.

## Discussion

Several physiological or environmental events can impact ER glycoproteostasis and glycoprotein trafficking along the secretory pathway. These conditions are known to induce temporary or chronic ER stress, which in turn leads to the activation of UPR pathways, with the aim of reducing the burden of ER stress, restoring glycoproteostasis or, if the ER stress condition persists, eventually triggering cell death. The molecular mechanisms controlling the fate of a given cell under ER stress are still largely to be elucidated. However, it is clear that the molecular cross-talk among different UPR arms and the signalling duration of each arm might be key determinants tipping the balance between survival and death. UPR arms have developed partially overlapping signalling pathways, rapidly activated in response to ER stress: studies on mutants of *bzip28*/*s1p* and *bzip60*/*ire1* clearly indicated a shared contribution of the two main arms of UPR in the activation of ER chaperones, e.g. BiP3, ERdj3B, CNX and PDI5 (Che et al., 2010; Ruberti et al., 2018). However, some of these genes are mainly controlled by bZIP60, as in the case of BiP3, while others mostly depend on bZIP28, e.g. ERdj3B (Ruberti et al., 2018). Furthermore, experiments using *ire1* mutants demonstrated a role for bZIP60 independent of its ribonuclease activity, implying that unspliced bZIP60 also plays a role in the recovery from ER stress (Henriquez-Valencia et al., 2015; Ruberti et al., 2018).

Most of the work done on UPR in plants has been carried out using drugs, such as tunicamycin (Tm) and DTT, impacting directly on glycoproteins folding. Under these stress conditions, the *bzip28/bzip60* double mutant showed reduced plant growth and reduced expression of BiP3 and CNX genes (Sun et al., 2013). Biotic stresses were also reported to induce ER stress and UPR, as in the case of the increased levels of *NbbZIP60* and BiP3 in *Nicotiana benthamiana* challenged by the pathogen *Pseudomonas cichori* (Tateda et al., 2008).

In contrast to stress induced by Tm, DTT or biotic stresses (the responses to which need the participation of both the bZIP28 and bZIP60 arms of UPR, with concomitant BiP3/CNX upregulation), salt/osmotic stress seems to be mainly controlled by the membrane associated bZIP17 TF, which requires proteolytic activation mediated by the Golgi subtilisin-like serine protease *At*S1P (Liu et al., 2007a). The involvement of bZIP60 in salt stress response is more controversial, since Henriquez-Valencia and co-workers (2015), did not detect the presence of *bZIP60* spliced transcripts nor chaperones (*CRT* or PDIL-1) under salt stress, even though *bZIP60* over-expression was reported to lead to a salt-tolerant phenotype (Fuijita et al., 2007). Although it is well known that Cd^2+^ treatment impacts negatively on the growth rate of *A. thaliana* Wt plants, with Cd-induced stress causing a significant reduction of primary root length and plant biomass (see also Fig. 1), a few reports have dealt so far with the effects of this source of stress on ER glycoproteostasis and activation of UPR. Our results clearly confirm that both chronic and temporary Cd^2+^ stress alter ER glycoproteostasis, with a concomitant upregulation of bZIP60 and bZIP28 TFs. Transient and harsher Cd^2+^ treatments result in a stronger induction of bZIP60 transcripts when compared with chronic stress, with the spliced form largely predominant over the unspliced transcript. bZIP28 transcription is induced equivalently in chronic and temporary Cd^2+^ stresses. Taken together, these results indicate that Cd^2+^ treatment activates mainly the bZIP60 arm, and therefore differs from other abiotic stresses.

Similarly to what is known about Tm- or DTT-induced ER stress, soon after Cd^2+^ stress is sensed, UPR signals, mainly through the bZIP60 arm, activate ERQC/ERAD to restore ER homeostasis and increase protein folding capacity. Indeed, BiP3, CNX, PDI5 and ERdj3B all show an expression trend similar to what observed for bZIP60, with concentration-(in the case of Cd^2+^ chronic stress) or time-dependent (temporary stress) effects (Lu and Christopher, 2008; Blanco-Herrera et al., 2015; Ruberti et al., 2018). A significant increase of both *AtbZIP60* and molecular chaperones transcripts is apparent even at the lowest Cd^2+^ concentration considered in this study (25 μM). These data confirm that UPR, under low stress conditions, is committed to reactivate ER glycoproteostasis and reduce the burden of ER accumulated unfolded glycoproteins. A more severe (50 μM Cd^2+^ during chronic stress) or longer transient stress (72 h) upregulates *NAC089*, activating PCD.

The *bzip28/bzip60* double mutant is more tolerant than Wt plants to Cd^2+^ stress, displaying significant recovery of primary root length and shoot fresh weight (Fig. 3). In this context our data confirm previous work from Xi and co-workers (2016), who reported for the first time that this mutant was less sensible to Cd^2+^ stress when compared with Wt plants challenged with 50 or 75 μM Cd^2+^. Transcription profiles of all the tested chaperones (*BiP3, ERdj3B, CNX, PDI5*) showed similar levels to those recorded in unstressed plants. The only exception was represented by the slight but significant increase of BiP3 under transient stress (48, 72 h). All these genes appear to be, at least in part, under the control of bZIP28/bZIP60, as already reported by others using knock-out mutant plants as the experimental model (Liu and Howell, 2016; Ruberti et al., 2018; Sun et al., 2021). In addition, an upregulation of them has been reported in plants overexpressing the active form of bZIP28 compared to Wt, even when grown in the control condition (Liu et al., 2007b).

Nevertheless, for the first time we can hypothesise that other TFs, i.e. bZIP17, may be involved in the slight upregulation of BiP3 under transient stress (Fig. 5C) as already reported for osmotic stresses (Liu et al., 2007a). Finally, the equivalent expression levels of *IRE1* observed in Wt and *bzip28/bzip60* mutant plants under Cd^2+^ stress clearly confirm previous reports, suggesting that this nuclease controls and regulates bZIP60 activation (Iwata and Koizumi, 2012; Mishiba et al., 2013; Li and Howell, 2021).

Overall, these data suggest that the increased Cd^2+^ resistance of the *bzip28/bzip60* mutant plants is due to a lower perception of ER stress and not to a lower amount of Cd^2+^ in tissues (Fig. 4; Fig. S1), highlighting a novel strategy to develop transgenic lines with improved tolerance and resilience.

Our data on BRI1, the brassinosteroid receptor which folds under ERQC control, confirm that BRI1 is an important regulator of plant growth under Cd^2+^ stress and the normal BRI1 glycoprotein levels observed in *bzip28/bzip60* could at least in part explain the better fitness of this mutant under Cd^2+^ stress. This is in agreement with a recent paper showing that brassinosteroid treatment increases Cd^2+^ tolerance (Della Rovere et al., 2022).

In a previous work, Blanco-Herrera and co-workers (2015) reported that the activation of UPR after Tm or DTT treatment induces the over-expression of *BiP3, PDI11*, and *UGGT* in *A. thaliana* plants. Our results indicate that Cd^2+^ stress also triggers UGGT but not *MNS4* overexpression. Of note is the downregulation of all these genes in the *bzip28/bzip60* mutant under normal growth conditions and under low Cd^2+^ stress conditions (25 μM), confirming that ERQC, ERAD and UPR are tightly inter-connected and inter-dependent mechanisms.

We did not observe significant differences in the overall glycoprotein folding capacity between Wt and the *bzip28/bzip60* mutant upon Cd^2+^ treatment. However, we did observe lower levels of EndoH sensitive glycoproteins in the Cd^2+^ stressed *bzip28/bzip60* mutant. Altogether these data point to a lower folding capacity for this mutant under Cd^2+^ stress conditions.

## Conclusions

The work sheds light on the *A. thaliana* response to Cd^2+^ stress. We show for the first time that ER stress and activation of UPR, due to a chronic or temporary Cd^2+^ treatment, proceeds mainly through the bZIP60 arm, although bZIP28 is also upregulated and contributes to the response. Our results also indicate that the high tolerance displayed by the *bzip28/bzip60* mutant to Cd^2+^ stress is at least in part due to an attenuated UPR-mediated response, with consequently reduced induction of ERQC chaperones under low Cd^2+^ stress conditions and PCD (*via* the activation of NAC089 TF) under harsher conditions. Higher BRI1 glycoprotein levels can also contribute to the better fitness observed in the double mutant.

The identification and characterisation of the finely tuned molecular mechanisms put in place by the plant cell to cope with ER stress triggered by heavy metals, together with the work carried out on metal transporters, will help to develop new varieties with potential applications in phytoremediation.

## Supplementary data

The following supplementary data are available at JXB online.

Fig. S1. Wt and *bzip28/bzip60* double mutant shown the same amount of Cd in their tissues. Fig. S2. Polyacrylamide gel of proteins extracted from Wt and *bzip28/bzip60* double mutant grown with or without CdCl_2_.

Fig. S3. Electrophoretic patterns of total proteins extracted from seedlings grown with or without CdCl_2_ and subjected to Endo H treatment.

Table S1. *A. thaliana* genes accession number

Table S2. List of primers used in qRT-PCR.

## Acknowledgements

We thank Prof. Mauro Marra (University of Rome “Tor Vergata”) for kindly providing seeds of the double mutant *bzip28/bzip60*. We also acknowledge Vittorio Falco for technical assistance.

## Author contributions

MDB and AG acquired the data. MDB, AG, PR and AS analyzed the results. DM and PP quantified Cd in *A. thaliana* plants. MDB, AG, PR and AS wrote the paper. All the authors critically revised the manuscript and approved the final version.

## Conflicts of interest

The authors declare no conflict of interest.

## Funding

This work was in part funded by the CNR-DiSBA excellence project Plant_EDEM.

## Data availability

The data underlying this article are available in the article and in its online supplementary material.

## Notes

### Competing Interest Statement

The authors have declared no competing interest.

